# Species identification of early colonial bone artefacts excavated from Pyrmont, Australia, by mass spectrometric identification of collagen peptides

**DOI:** 10.1101/2022.05.13.491741

**Authors:** Dylan H. Multari, Geraldine J. Sullivan, Mary Hartley, Ronika K. Power, Paul A. Haynes

**Affiliations:** School of Natural Sciences, Macquarie University, Sydney, Australia; Department of History and Archaeology, Macquarie University, Sydney, Australia; Biomolecular Discovery Research Centre, Macquarie University, Sydney, Australia

**Keywords:** Colonial-era Australia, tandem Mass Spectrometry, Paleoproteomics, Zooarchaeology

## Abstract

Zooarchaeology by Mass Spectrometry (ZooMS) is a rapidly developing and increasingly utilised peptide mass fingerprinting (PMF) technique that analyses Collagen 1A1 and 1A2 marker peptides for the genus- or species-level identification of fragmentary bones in the archaeological record. Traditionally, this analysis is performed using matrix-assisted laser desorption/ionisation time-of-flight mass spectrometry (MALDI-ToF-MS) to identify characteristic m/z values of known marker peptides. Here we present data on the application of a modified ZooMS approach, using nanoflow liquid chromatography - tandem mass spectrometry proteomics, to the analysis of a collection of six early colonial Australian (early to mid-19^th^ Century CE) worked bone artefacts, believed to be mostly knife handles, excavated from a site in Pyrmont, Sydney, Australia in 2017. We were successfully able to identify characteristic marker peptides for bovine COL1A1 and COL1A2 in all six bone artefacts.

## 1. Introduction

Zooarchaeology by Mass Spectrometry (ZooMS) is an established peptide mass fingerprinting (PMF) technique that commonly utilises matrix-assisted laser desorption/ionisation – tandem time of flight mass spectrometry (MALDI-ToF/ToF) to identify genus- or species-specific COL1A1 or COL1A2 peptide markers from fragmentary bones in the archaeological record (Buckley, et al., 2009, McGrath, et al., 2019, Brown, et al., 2021, Wang, et al., 2021). The utility of this collagen PMF technique lies in its cost-effectiveness, reduced sample requirement, and the ability of proteins, such as collagens, to survive across near-geological time periods and harsh environments, where ancient DNA analysis would likely not be feasible (Demarchi, et al., 2016, Wang, et al., 2021). This, coupled with the intrinsic connection between the DNA sequence and protein sequence in an organism (commonly referred to as the Central Dogma of molecular biology), is what allows ZooMS to be a robust alternative to aDNA analysis in these contexts.

ZooMS has been applied to a wide variety of samples from extensive geographical and temporal contexts and has been previously reviewed in, for example: Hendy (2021), and Schroeter, et al. (2022). This technique has been applied to a range of archaeological problems including resolving the evolutionary and phylogenetic history of Darwin’s South American ungulates (Welker, et al., 2015), determining the phylogenetic relationship between a series of extinct giant ground sloths (Buckley, et al., 2015), and distinguishing between mammoth and mastodon fossils (Buckley, et al., 2011). However, very few studies exist in the literature of the application of ZooMS analysis to faunal remains from an Australian context, with none pertaining to relatively recent colonial-era history (Buckley, et al., 2017, Peters, et al., 2021). Here we present the results of the analysis of a collection of six early colonial Australian (c. 1830) bone artefacts excavated from a site in Pyrmont, Sydney, Australia, using a modified ZooMS approach. We compared three established collagen extraction techniques (Wang, et al., 2021) on small aliquots of bone powder, followed by nanoflow high-pressure liquid chromatography – tandem mass spectrometry (NanoLC-MS/MS), for the taxonomic identification of collagen peptide markers extracted from the bone artefacts.

## 2. Materials and Methods

### 2.1 Site context

In the early colonisation of Sydney, the Pyrmont Peninsula (whose traditional name was “Pirrama”) was the home of the Cadigal/Gadigal people who were later displaced in the 1830s as the colony began to spread and industrialise (Fitzgerald and Golder, 1994). During this period of development, the region became a ‘working-class’ district which housed a large population of poorer settlers, as well as a number of industries including abattoirs and breweries (Fitzgerald and Golder, 1994). In 2017, EMM Consulting was commissioned to perform an archaeological assessment and subsequent excavation of the site for a proposed redevelopment of the land (EMM Consulting Pty Limited, 2020). Figure 1 shows the excavation site in a modern regional context, illustrating its position in the centre of what is now a largely developed urban area in the City of Sydney. The excavation site was dated to approximately the early to mid-19^th^ Century based on archival evidence. For example, Figure 2 is a historical map of the Pyrmont Peninsula which shows the development and subdivision plans proposed by Edward Macarthur in 1836 after inheriting the land from his late father. The excavation site is marked with a purple box in Figure 2, and a more detailed excavation site map of 63, 65A, and 69-71 Harris Street and 14-16 Mount Street, Pyrmont, subdivided into the respective excavation contexts, can be seen in Figure 3. A large collection of animal bones representing various taxa were found during the excavation of this site, including a collection of worked bone artefacts (Figure 4).

**Figure 1:**
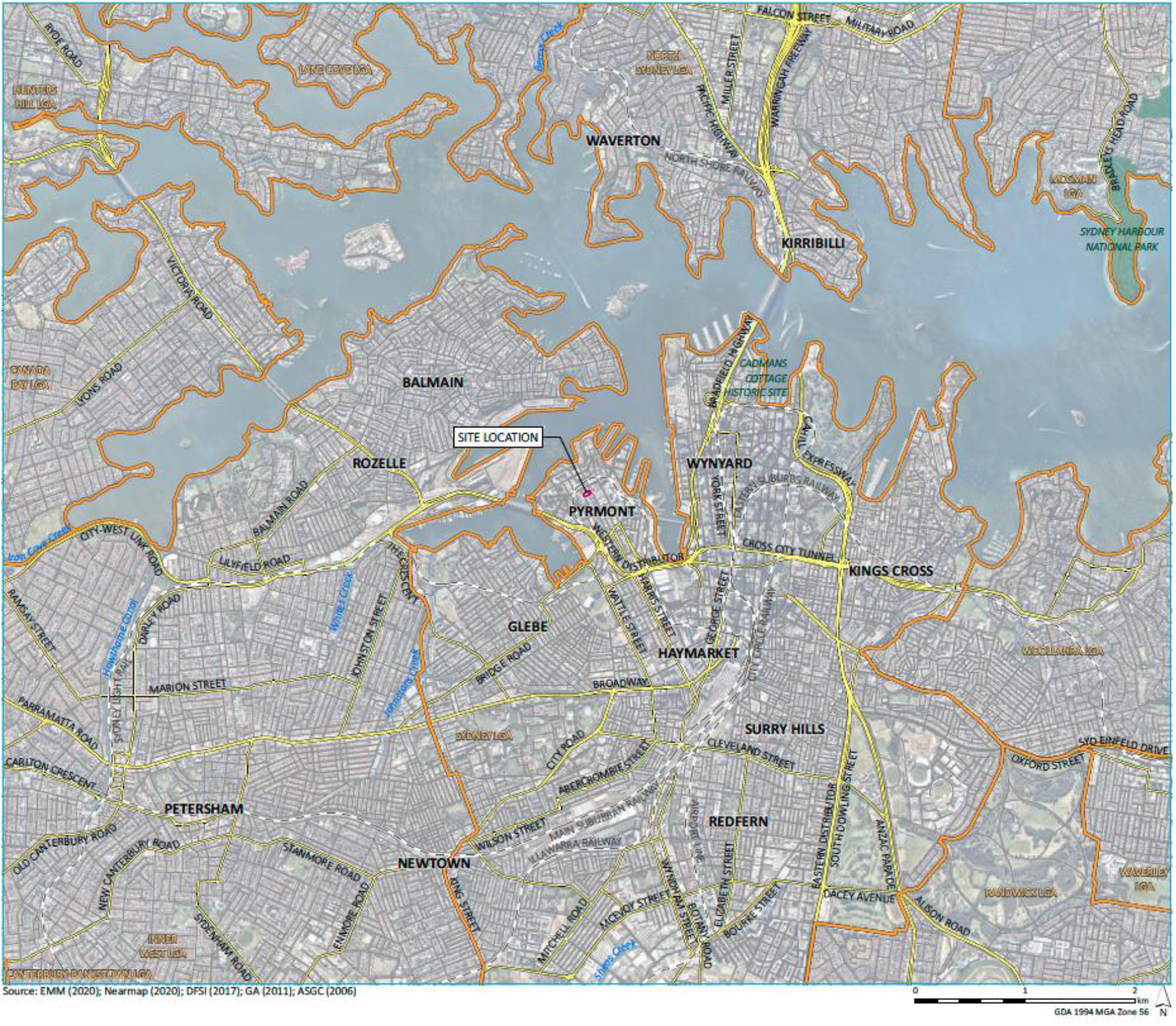
Map of Sydney Harbour region showing the subject site in a regional context. Excavation site is denoted by a purple square in the Pyrmont area. Map courtesy of Pamela Kottaras from EMM Consulting. Image used with express permission of EMM Consulting.

**Figure 2:**
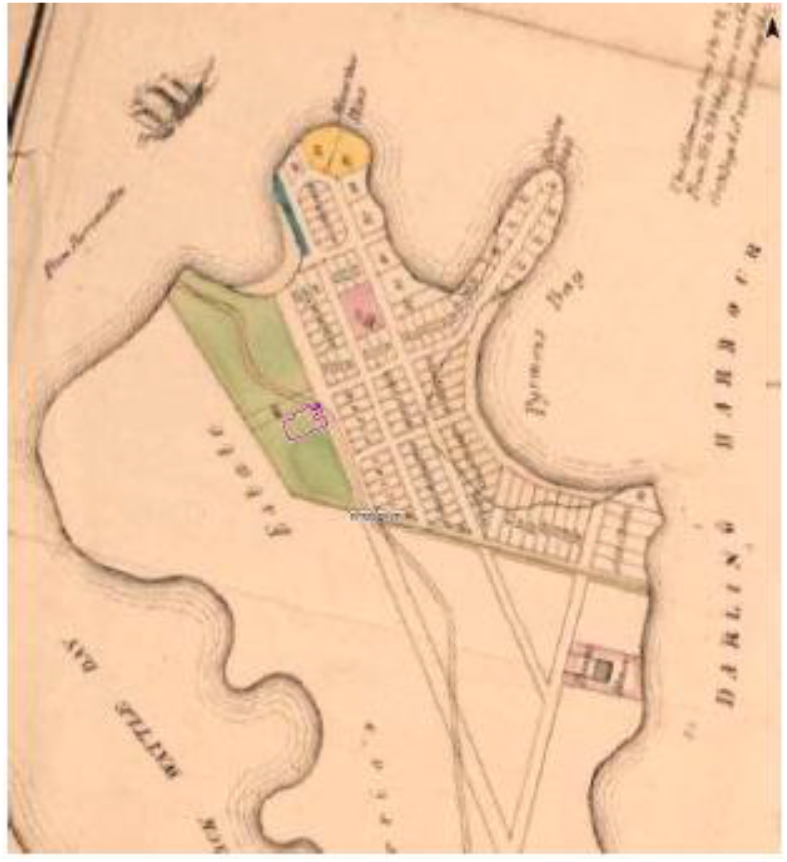
Edward Macarthur’s plan for the subdivision and development of the Pyrmont Peninsula, Sydney, Australia, 1836. Image from the National Library of Australia, Call. No. MAP T 1551. Site of excavation is outlined in purple.

**Figure 3:**
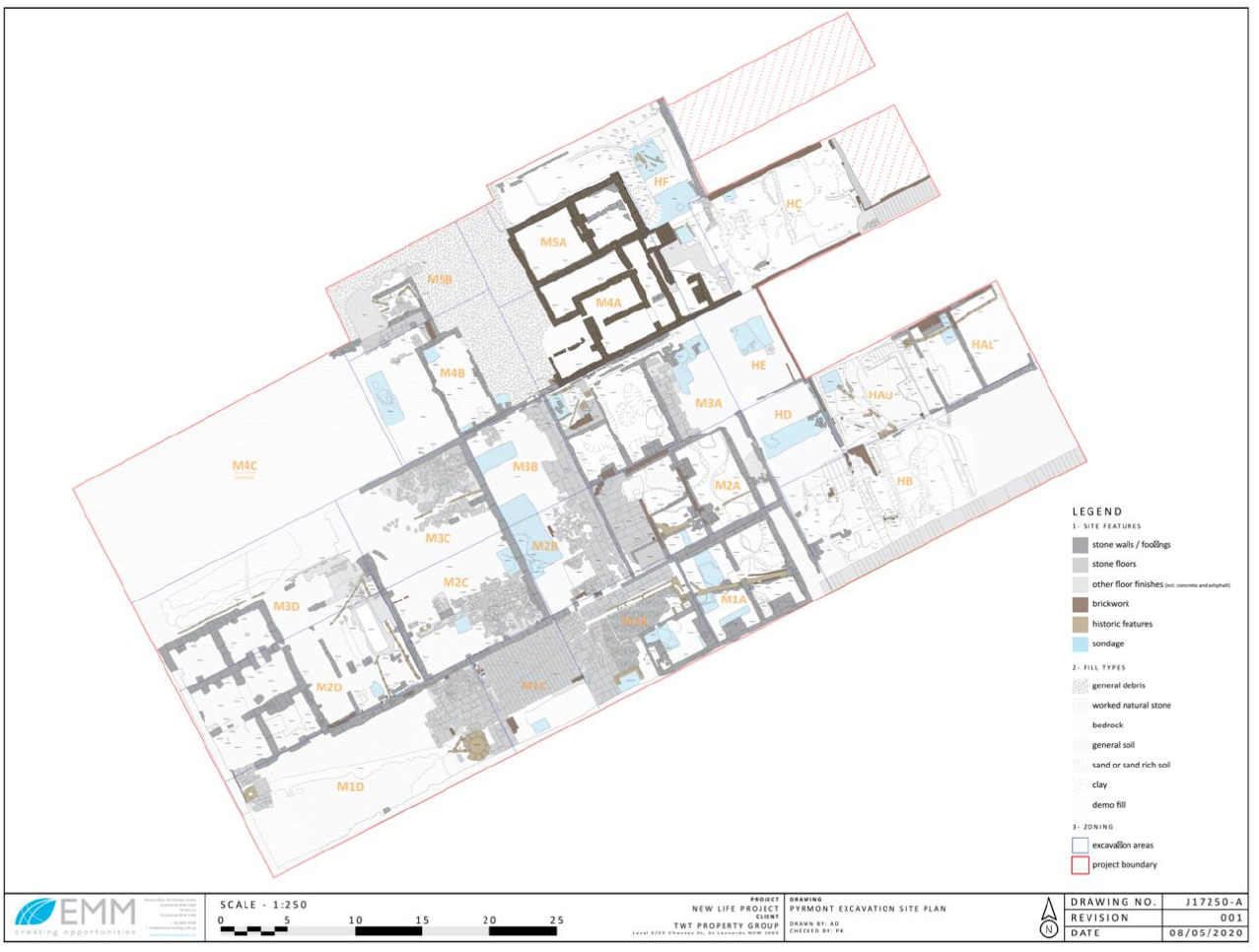
Excavation Site Map of 63, 65A & 69-71 Harris Street and 14-16 Mount Street, Pyrmont, Sydney, Australia. Image courtesy of Pamela Kottaras from EMM Consulting. Image used with express permission of EMM Consulting.

**Figure 4:**
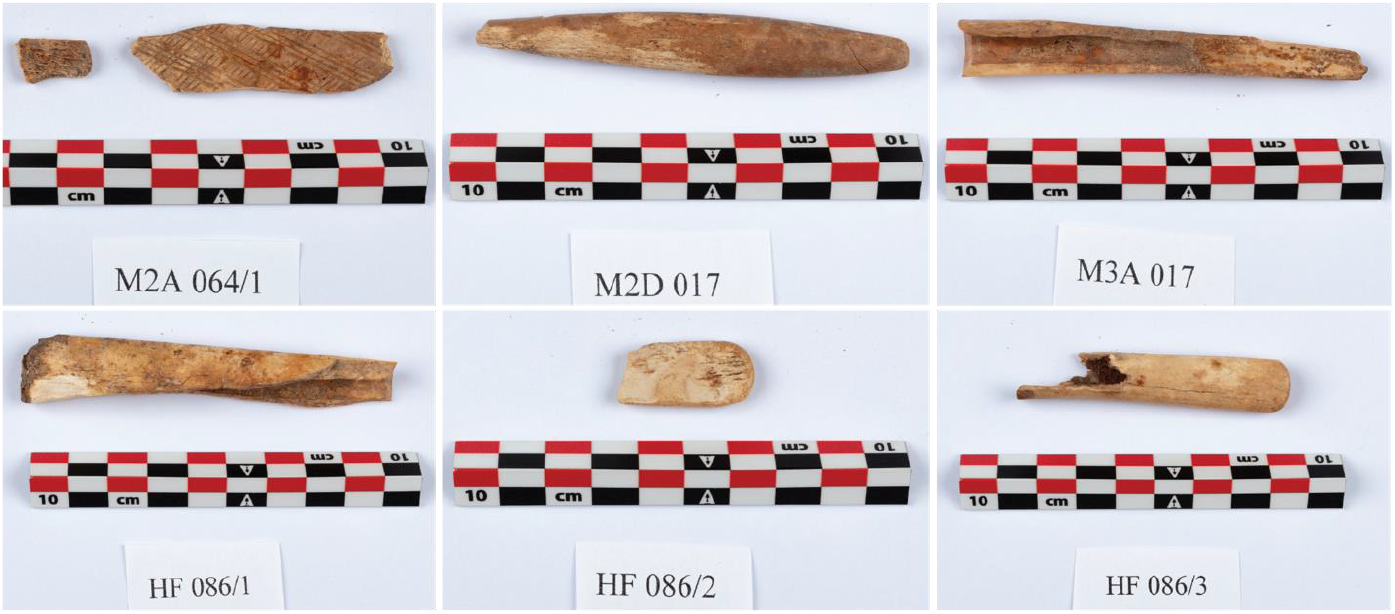
Collection of bone artefacts from 63, 65A & 69-71 Harris Street and 14-16 Mount Street, Pyrmont excavation site. (Left to right) M2A 064.1; M2D 017; M3A 017; HF 086.1; HF 086.2; and HF 086.3. Scale bar is 10 cm. Images courtesy of Michael Rampe.

### 2.2 Materials

A collection of six bone artefacts were found during the excavation at 63, 65A & 69-71 Harris Street and 14-16 Mount Street, Pyrmont (Figure 4). Initial post-excavation cataloguing of these artefacts listed them as knife handles. The length of the artefacts ranges from approximately 2 cm to 10 cm, with an average width of between 1 to 2 cm. Due to their worked nature, morphological identification as typically performed in osteological zooarchaeology was not possible. Artefacts M3A 017, HF 086.1, and HF 086.3 had hollowed-out interiors, most likely the modified medullary cavities of metapodial bones (Paral, et al., 2004, Unwin, 2018), while the remaining handles were solid pieces of bone. It is possible that the artefact M2D 017 belonged to a toothbrush owing to the tapered shape of the bone, however this is hard to discern as it was not fully intact upon excavation. Interestingly, M2A 064.1 displayed a decorative hatching pattern inscribed onto the cortical surface of the bone that was not evident on any other bone piece excavated from this site. This artefact was identified as a rib bone, and based on the thickness and decoration, it is likely to have been an inlay rather than a knife handle itself.

### 2.3 Sampling of bone powder

Special care to avoid contamination was taken in the sample handling of the bone artefacts and in the subsequent laboratory analysis, as suggested for ancient protein experiments (Hendy, et al., 2018). Namely, this involved the use of nitrile gloves and face mask, sterilised equipment and workspaces, freshly prepared reagents, and processing of analytical and procedural blanks in addition to samples. Laboratory workspace was covered in aluminium foil prior to sampling to collect any loose powder. Approximately 15 mg of bone powder was collected from each artefact using a Dremel sanding tool into a fresh weigh boat. The Dremel sanding tool and laboratory space were thoroughly cleaned with 80% EtOH (v/v) after each sample to minimise potential cross-contamination. Bone powder was washed twice briefly in 100 μL of 50 mM NH_4_HCO_3_ to remove soluble surface contaminants.

### 2.4 Collagen Extraction

To maximise the amount of potential marker peptide identifications, three different collagen extraction protocols which have been previously reported in the literature (Wang, et al., 2021) were applied to the collected bone powder. Briefly, these are the Acid Soluble (AC), Acid Insoluble (AI), and the non-destructive AmBiC (AM) protocols. Sampling was performed twice for each bone artefact to collect enough bone powder to perform each of the protocols.

#### 2.4.1 Acid Soluble Protocol

The portions of bone powder designated for the acid-based protocols were demineralised at 4°C in 500 μL of 600 mM HCl until there were no visible signs of reaction. Samples were centrifuged briefly at 14,500 rpm for one minute, then the acid supernatant was transferred to a 30 kDa centrifuge spin filter (Microcon-30 kDa Centrifugal Filter Unit with Ultracel-30 membrane, Merck, Australia; Product ID: MRCF0R030) prior to further centrifugation at 3,700 rpm for approximately 30 minutes until liquid had passed through the filter membrane. Flow-through was discarded, and 500 μL of 50 mM NH_4_HCO_3_ was added to the spin filter unit. Centrifugation was repeated at 3,700 rpm until the liquid had passed through, and flow-through was similarly discarded. An aliquot of 100 μL of 50 mM NH_4_HCO_3_ was added to the filter unit and collagen extracts were resuspended via gentle pipetting before transferring to a fresh 1.5 mL microcentrifuge tube for subsequent trypsin digestion.

#### 2.4.2 Acid Insoluble Protocol

HCl-treated bone powder was re-equilibrated via washing with 100 μl of 50 mM NH_4_HCO_3_ briefly. Bone powder was centrifuged briefly at 14,500 rpm and NH_4_HCO_3_ was removed, then replaced with a fresh 100 μL aliquot of 50 mM NH_4_HCO_3_. These samples were then gelatinised at 65°C for 1 hour. After gelatinisation, samples were spun briefly at 14,500 rpm and the supernatant was transferred to a fresh 1.5 mL microcentrifuge tube for subsequent trypsin digestion.

#### 2.4.3 AmBiC Protocol

The remaining untreated bone powder sample was washed briefly with 100 μL 50 mM NH_4_HCO_3_, then gelatinised in a fresh aliquot of 100 μL 50 mM NH_4_HCO_3_ at 65°C for 1 hour as previously described in the Acid Insoluble protocol. Samples were then spun briefly at 14,500 rpm and supernatant was transferred to a fresh 1.5 mL microcentrifuge tube for trypsin digestion.

### 2.5 Trypsin in-solution digestion and C18 Zip-tip purification of resultant peptides

Collagen extracts from each sample preparation method were digested in solution with 0.4 μg/μL trypsin (Sequencing Grade Modified Porcine Trypsin, Promega) in 100 mM NH_4_HCO_3_ overnight at 37°C. Peptide clean-up was performed using OMIX C18 100 μL tips (Agilent Technologies, Product ID: A57003100). C18 tips were equilibrated by washing twice with 50 μL Conditioning Solution (80% Acetonitrile (ACN)/ 0.1% Trifluoroacetic acid (TFA)), then twice with 50 μL Washing Solution (2% ACN/ 0.1% TFA). Peptides were loaded onto the C18 resin via pipetting back and forth at least 10 times, then washed twice with Washing Solution, and eluted by pipetting 50 μL Elution Solution (60% ACN/ 0.1% TFA) at least 10 times to ensure thorough elution into a fresh 1.5 mL microcentrifuge tube. This was repeated for each sample using fresh Zip-tips and reagents. Purified samples were vacuum centrifuged to dryness and reconstituted in 10 μL 1% Formic Acid (FA) for subsequent LC-MS/MS analysis.

### 2.6 NanoLC-MS/MS of peptides

NanoLC-MS/MS was performed on a Thermo Q-Exactive orbitrap mass spectrometer coupled to a Thermo Easy-nLC1000 liquid chromatography system as previously described (Muralidharan, et al., 2012, Mirzaei, et al., 2014, Multari, et al., 2022), with minor modifications. Briefly, collagen peptide digests were loaded onto a 75 μm ID × 100 mm length column packed with C18 HALO resin (2.7 μm bead size, 160 Å pore size) for reversed-phase chromatographic separation. A linear gradient of 1-50% solvent A (0.1% FA) to solvent B (99.9% ACN/ 0.1% FA) was developed over 60 minutes. Data-dependent acquisition was configured to automatically switch from Orbitrap MS to MS/MS mode, and spectra were acquired from a m/z range of 350-1600 amu with a resolution of 35,000 and an isolation window of 3.0 m/z. The top ten most abundant ions were subsequently selected for higher energy collisional dissociation (HCD) fragmentation at 30% normalised HCD collision energy, with a dynamic exclusion of target ions set for 20 seconds. Fragmentation ions were detected in the orbitrap at a resolution of 17,500. Analytical blanks of 70% MeOH (v/v) were run between samples to minimise potential system carryover of abundant peptides. Raw mass spectrometric data were deposited to the ProteomeXchange Consortium via the PRIDE (Perez-Riverol, et al., 2019) partner repository.

### 2.7 Protein identification using X! Tandem

Raw mass spectral files were converted to mzXML format using MSConvertGUI (Kessner, et al., 2008, Chambers, et al., 2012), and peptide-to-spectrum matching was performed using the X! Tandem algorithm under the Global Proteome Machine (GPM) user interface software (version 3.0, https://www.thegpm.org/) (Craig and Beavis, 2004, Craig, et al., 2004). Converted mzXML files were searched against the curated SwissProt protein database (downloaded April 2020, 562,895 proteins) supplemented with additional COL1A1 and COL1A2 sequences for species of interest including all available mammalian COL1A1 and COL1A2 sequences from the NCBI protein database, previously reported experimentally-determined marsupial peptide markers (Buckley, et al., 2017, Peters, et al., 2021), and the Common Repository of Adventitious Proteins (cRAP) database (Shin, et al., 2019). Searching was performed with the following specified parameters: Orbitrap instrument method (±20 ppm parent ion mass tolerance), fragment ion mass tolerance ±0.1 Da, trypsin digestion with allowance for two missed cleavages, minimum and maximum peptide sequence lengths of 6 and 50 amino acids, respectively, and up to 4+ charge states allowed. No fixed modifications were specified, however variable modifications were as follows: oxidation of methionine and tryptophan, hydroxylation of proline, and deamidation of glutamine and asparagine. Spectra were searched against a decoy reversed sequence database for assessment of false discovery rates (FDR). Collated GPM peptide outputs can be found in Supplementary Data S1.

### 2.8 Data visualisation and analysis

Deamidation assessment was performed using the R coding language (version 3.6.3) (R Core Team, 2020) operating in RStudio (version 1.3.1073) using an in-house script as previously described (Multari, et al., 2022). GPM peptide output files were assessed for the presence of identified peptide markers, which were then extracted into a separate Excel spreadsheet containing details about sample, method, ZooMS marker region, peptide sequence, and reported species of origin (Supplementary Data S2) for subsequent data visualisation (Figure 5) and analysis (Table 2) in RStudio. Each unique α2 peptide sequence was collated as a single sequence for each sample into a FASTA file containing a series of COL1A2 sequences for common species and select Australian native animals for sequence alignment using the *msa* package (version 1.18.0) (Bodenhofer, et al., 2015), and pairwise sequence identity distance alignment using a Fitch matrix (Fitch, 1966) in the R package *seqinr* (version 4.2-8) (Charif and Lobry, 2007). COL1A2 sequence alignments are supplied in Supplementary Data S3.

## 3. Results

### 3.1 Identification of ZooMS Collagen peptide markers

Collagen peptide markers for ZooMS analysis have been previously reported, along with a standardised nomenclature for reporting the identification of such peptides (Brown, et al., 2021). Table 1 shows the number of times each reported ZooMS marker region peptide was identified for each sample separated by sample preparation method (AC, AI, or AM). Figure 5 illustrates the summed distribution of marker region identifications across sample preparation methods. The α2 757 peptide region was not identified in any of the samples across all three collagen extraction methods.

**Table 1.** Summary table of ZooMS Peptide Markers (Brown, et al., 2021) identified via LC-MS/MS using the three collagen extraction techniques. Observation count also includes peptides with one missed tryptic cleavage that contain the entire ZooMS peptide marker region.

| Sample | ZooMS Peptide Marker (number of observations) |  |  |  |  |  |  |  |  |
| --- | --- | --- | --- | --- | --- | --- | --- | --- | --- |
| | $\alpha 1$<br>508 | $\alpha 1$<br>586 | $\alpha 2$<br>292 | $\alpha 2$<br>454 | $\alpha 2$<br>484 | $\alpha 2$<br>502 | $\alpha 2$<br>757 | $\alpha 2$<br>793 | $\alpha 2$<br>978 |
| AC |  |  |  |  |  |  |  |  |  |
| M2A 064.1 | 14 | 12 | 2 | 2 | 5 | 3 | - | 8 | 5 |
| M2D 017 | 11 | 14 | 2 | 2 | 6 | 9 | - | 5 | 6 |
| M3A 017 | 6 | 33 | 2 | 2 | 20 | 6 | - | 77 | 11 |
| HF 086.1 | 6 | 27 | 2 | - | 10 | 6 | - | 62 | 11 |
| HF 086.2 | 1 | 29 | 7 | 1 | 6 | 5 | - | 22 | 10 |
| HF 086.3 | 3 | 19 | 2 | 2 | 3 | 10 | - | 8 | 8 |
| AI |  |  |  |  |  |  |  |  |  |
| M2A 064.1 | 12 | 57 | 2 | 4 | 5 | 7 | - | 14 | 7 |
| M2D 017 | 5 | 6 | 9 | 1 | 4 | 8 | - | 5 | 30 |
| M3A 017 | 3 | 62 | 3 | 5 | 14 | 10 | - | 94 | 15 |
| HF 086.1 | 2 | 2 | - | - | 1 | 4 | - | 5 | 3 |
| HF 086.2 | - | 25 | 5 | 6 | 7 | 1 | - | 7 | 14 |
| HF 086.3 | 3 | 6 | 2 | 1 | 4 | 11 | - | 6 | 7 |
| AM |  |  |  |  |  |  |  |  |  |
| M2A 064.1 | 21 | 15 | 4 | 2 | 7 | 5 | - | 11 | 7 |
| M2D 017 | 9 | 12 | 3 | 4 | 6 | 12 | - | 6 | 9 |
| M3A 017 | 2 | 32 | 5 | 1 | 9 | 7 | - | 29 | 10 |
| HF 086.1 | 6 | 18 | 5 | 3 | 36 | 7 | - | 86 | 11 |
| HF 086.2 | 1 | 22 | 5 | 1 | 11 | 2 | - | 19 | 13 |
| HF 086.3 | 7 | 3 | 1 | 2 | 2 | 13 | - | 3 | 5 |

**Figure 5:**
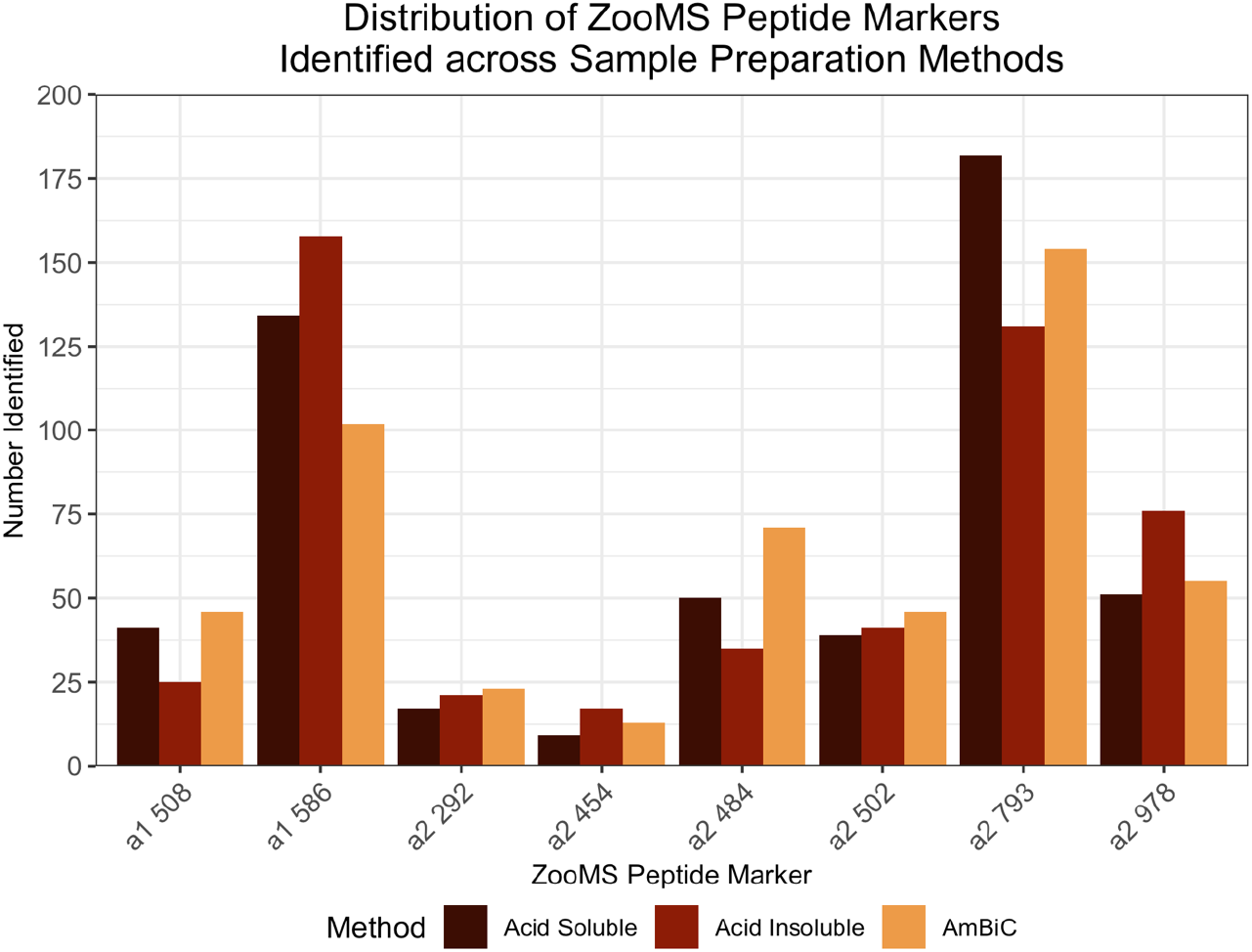
Distribution of ZooMS Peptide Markers identified across the three sample preparation methods. Figure generated from collated ZooMS peptides (Supplementary Data S2) using R (version 3.6.3) in RStudio (version 1.3.1073) (R Core Team, 2020).

### 3.2 Species identification of bone artefacts

The species identification of the bone artefacts was performed by analysing the peptide sequence data for the COL1A2 peptide marker regions (Table 1) identified via LC-MS/MS. The sequences obtained from each artefact were collated into a FASTA file containing a series of COL1A2 sequences for common species and select Australian native animals under a header consisting of the sample name. Table 2 highlights the pairwise distance alignment of the protein sequence identities using a Fitch matrix, where values from 0.0 to 1.0 represent percentage of variation in sequence identity. Values of 0.0 represent a direct correlation. Table 2 is reported at 1 significant figure, while raw un-rounded values are included in Supplementary Data S4. The pairwise analysis shows a direct correlation between the peptide sequence identity of each of the artefacts and between the COL1A2 sequence for *Bos taurus*.

**Table 2:** Pairwise distance matrix of aligned protein sequences for identified COL1A2 peptides from the bone artefacts and COL1A2 sequences for various common species and select Australian native animals. Distance alignment matrix was generated using a Fitch matrix (Fitch, 1966) in R (version 3.6.3) using the ‘seqinr’ package (4.2-8) (Charif and Lobry, 2007). Values rounded to 1 significant figure. Zero values are bolded for ease of reading. Raw values are included in Supplementary Data S4.

|  | HF 086.1 | HF 086.2 | HF 086.3 | M2A 064.1 | M2D 017 | M3A 017 | *B. taurus* | *C. hircus* | *O. aries* | *P. catodon* | *S. scrofa domesticus* | *C. lupus dingo* | *C. lupus familiaris* | *F. catus* | *E. caballus* | *R. rattus* | *M. musculus* | *H. sapiens* | *G. gorilla gorilla* | *P. troglodytes* | *T. aculeatus* | *O. anatinus* | *V. ursinus* | *P. cinereus* | *T. vulpecula* | *S. harrisi* |
| --- | --- | --- | --- | --- | --- | --- | --- | --- | --- | --- | --- | --- | --- | --- | --- | --- | --- | --- | --- | --- | --- | --- | --- | --- | --- | --- |
| HF 086.1 | **0.0** | **0.0** | **0.0** | **0.0** | **0.0** | **0.0** | **0.0** | 0.1 | 0.1 | 0.3 | 0.3 | 0.3 | 0.3 | 0.4 | 0.3 | 0.4 | 0.4 | 0.4 | 0.4 | 0.4 | 0.5 | 0.4 | 0.4 | 0.4 | 0.4 | 0.4 |
| HF 086.2 | **0.0** | **0.0** | **0.0** | **0.0** | **0.0** | **0.0** | **0.0** | 0.1 | 0.1 | 0.3 | 0.3 | 0.3 | 0.3 | 0.4 | 0.3 | 0.4 | 0.4 | 0.4 | 0.4 | 0.4 | 0.5 | 0.4 | 0.4 | 0.4 | 0.4 | 0.4 |
| HF 086.3 | **0.0** | **0.0** | **0.0** | **0.0** | **0.0** | **0.0** | **0.0** | 0.1 | 0.1 | 0.3 | 0.3 | 0.3 | 0.3 | 0.4 | 0.3 | 0.4 | 0.4 | 0.4 | 0.4 | 0.4 | 0.5 | 0.4 | 0.4 | 0.4 | 0.4 | 0.4 |
| M2A 064.1 | **0.0** | **0.0** | **0.0** | **0.0** | **0.0** | **0.0** | **0.0** | 0.1 | 0.1 | 0.3 | 0.3 | 0.3 | 0.3 | 0.4 | 0.3 | 0.4 | 0.4 | 0.4 | 0.4 | 0.4 | 0.5 | 0.4 | 0.4 | 0.4 | 0.4 | 0.4 |
| M2D 017 | **0.0** | **0.0** | **0.0** | **0.0** | **0.0** | **0.0** | **0.0** | 0.1 | 0.1 | 0.3 | 0.3 | 0.3 | 0.3 | 0.4 | 0.3 | 0.4 | 0.4 | 0.4 | 0.4 | 0.4 | 0.5 | 0.4 | 0.4 | 0.4 | 0.4 | 0.4 |
| M3A 017 | **0.0** | **0.0** | **0.0** | **0.0** | **0.0** | **0.0** | **0.0** | 0.1 | 0.1 | 0.3 | 0.3 | 0.3 | 0.3 | 0.4 | 0.3 | 0.4 | 0.4 | 0.4 | 0.4 | 0.4 | 0.5 | 0.4 | 0.4 | 0.4 | 0.4 | 0.4 |
| *Bos taurus* | **0.0** | **0.0** | **0.0** | **0.0** | **0.0** | **0.0** | **0.0** | 0.1 | 0.1 | 0.2 | 0.2 | 0.2 | 0.2 | 0.2 | 0.2 | 0.3 | 0.3 | 0.3 | 0.3 | 0.3 | 0.4 | 0.4 | 0.4 | 0.4 | 0.4 | 0.4 |
| *Capra hircus* | 0.1 | 0.1 | 0.1 | 0.1 | 0.1 | 0.1 | 0.1 | **0.0** | **0.0** | 0.2 | 0.2 | 0.2 | 0.2 | 0.2 | 0.2 | 0.3 | 0.3 | 0.3 | 0.3 | 0.3 | 0.4 | 0.4 | 0.4 | 0.4 | 0.4 | 0.4 |
| *Ovis aries* | 0.1 | 0.1 | 0.1 | 0.1 | 0.1 | 0.1 | 0.1 | **0.0** | **0.0** | 0.2 | 0.2 | 0.2 | 0.2 | 0.2 | 0.2 | 0.3 | 0.3 | 0.3 | 0.3 | 0.3 | 0.4 | 0.4 | 0.4 | 0.4 | 0.4 | 0.4 |
| *Physeter catodon* | 0.3 | 0.3 | 0.3 | 0.3 | 0.3 | 0.3 | 0.2 | 0.2 | 0.2 | **0.0** | 0.2 | 0.3 | 0.3 | 0.3 | 0.3 | 0.3 | 0.3 | 0.3 | 0.3 | 0.3 | 0.4 | 0.4 | 0.4 | 0.4 | 0.4 | 0.4 |
| *Sus scrofa domesticus* | 0.3 | 0.3 | 0.3 | 0.3 | 0.3 | 0.3 | 0.2 | 0.2 | 0.2 | 0.2 | **0.0** | 0.2 | 0.2 | 0.2 | 0.2 | 0.3 | 0.3 | 0.2 | 0.3 | 0.3 | 0.4 | 0.4 | 0.4 | 0.4 | 0.4 | 0.4 |
| *Canis lupus dingo* | 0.3 | 0.3 | 0.3 | 0.3 | 0.3 | 0.3 | 0.2 | 0.2 | 0.2 | 0.3 | 0.2 | **0.0** | **0.0** | 0.1 | 0.2 | 0.3 | 0.3 | 0.2 | 0.2 | 0.2 | 0.4 | 0.4 | 0.4 | 0.4 | 0.4 | 0.4 |
| *Canis lupus familiaris* | 0.3 | 0.3 | 0.3 | 0.3 | 0.3 | 0.3 | 0.2 | 0.2 | 0.2 | 0.3 | 0.2 | **0.0** | **0.0** | 0.1 | 0.2 | 0.3 | 0.3 | 0.2 | 0.2 | 0.2 | 0.4 | 0.4 | 0.4 | 0.4 | 0.4 | 0.4 |
| *Felis catus* | 0.4 | 0.4 | 0.4 | 0.4 | 0.4 | 0.4 | 0.2 | 0.2 | 0.2 | 0.3 | 0.2 | 0.1 | 0.1 | **0.0** | 0.2 | 0.3 | 0.3 | 0.2 | 0.2 | 0.2 | 0.4 | 0.4 | 0.4 | 0.4 | 0.4 | 0.4 |
| *Equus caballus* | 0.3 | 0.3 | 0.3 | 0.3 | 0.3 | 0.3 | 0.2 | 0.2 | 0.2 | 0.3 | 0.2 | 0.2 | 0.2 | 0.2 | **0.0** | 0.3 | 0.3 | 0.3 | 0.3 | 0.3 | 0.4 | 0.4 | 0.4 | 0.4 | 0.4 | 0.4 |
| *Rattus rattus* | 0.4 | 0.4 | 0.4 | 0.4 | 0.4 | 0.4 | 0.3 | 0.3 | 0.3 | 0.3 | 0.3 | 0.3 | 0.3 | 0.3 | 0.3 | **0.0** | 0.2 | 0.3 | 0.3 | 0.3 | 0.4 | 0.4 | 0.4 | 0.4 | 0.4 | 0.4 |
| *Mus musculus* | 0.4 | 0.4 | 0.4 | 0.4 | 0.4 | 0.4 | 0.3 | 0.3 | 0.3 | 0.3 | 0.3 | 0.3 | 0.3 | 0.3 | 0.3 | 0.2 | **0.0** | 0.3 | 0.3 | 0.3 | 0.4 | 0.4 | 0.4 | 0.4 | 0.4 | 0.4 |
| *Homo sapiens* | 0.4 | 0.4 | 0.4 | 0.4 | 0.4 | 0.4 | 0.3 | 0.3 | 0.3 | 0.3 | 0.2 | 0.2 | 0.2 | 0.2 | 0.3 | 0.3 | 0.3 | **0.0** | 0.1 | **0.0** | 0.4 | 0.4 | 0.4 | 0.4 | 0.4 | 0.4 |
| *Gorilla gorilla gorilla* | 0.4 | 0.4 | 0.4 | 0.4 | 0.4 | 0.4 | 0.3 | 0.3 | 0.3 | 0.3 | 0.3 | 0.2 | 0.2 | 0.2 | 0.3 | 0.3 | 0.3 | 0.1 | **0.0** | 0.1 | 0.4 | 0.4 | 0.4 | 0.4 | 0.4 | 0.4 |
| *Pan troglodytes* | 0.4 | 0.4 | 0.4 | 0.4 | 0.4 | 0.4 | 0.3 | 0.3 | 0.3 | 0.3 | 0.3 | 0.2 | 0.2 | 0.2 | 0.3 | 0.3 | 0.3 | **0.0** | 0.1 | **0.0** | 0.4 | 0.4 | 0.4 | 0.4 | 0.4 | 0.4 |
| *Tachyglossus aculeatus* | 0.5 | 0.5 | 0.5 | 0.5 | 0.5 | 0.5 | 0.4 | 0.4 | 0.4 | 0.4 | 0.4 | 0.4 | 0.4 | 0.4 | 0.4 | 0.4 | 0.4 | 0.4 | 0.4 | 0.4 | **0.0** | 0.2 | 0.4 | 0.4 | 0.4 | 0.4 |
| *Ornithorhynchus anatinus* | 0.4 | 0.4 | 0.4 | 0.4 | 0.4 | 0.4 | 0.4 | 0.4 | 0.4 | 0.4 | 0.4 | 0.4 | 0.4 | 0.4 | 0.4 | 0.4 | 0.4 | 0.4 | 0.4 | 0.4 | 0.2 | **0.0** | 0.4 | 0.4 | 0.4 | 0.4 |
| *Vombatus ursinus* | 0.4 | 0.4 | 0.4 | 0.4 | 0.4 | 0.4 | 0.4 | 0.4 | 0.4 | 0.4 | 0.4 | 0.4 | 0.4 | 0.4 | 0.4 | 0.4 | 0.4 | 0.4 | 0.4 | 0.4 | 0.4 | 0.4 | **0.0** | 0.2 | 0.2 | 0.2 |
| *Phascolarctos cinereus* | 0.4 | 0.4 | 0.4 | 0.4 | 0.4 | 0.4 | 0.4 | 0.4 | 0.4 | 0.4 | 0.4 | 0.4 | 0.4 | 0.4 | 0.4 | 0.4 | 0.4 | 0.4 | 0.4 | 0.4 | 0.4 | 0.4 | 0.2 | **0.0** | 0.2 | 0.2 |
| *Trichosurus vulpecula* | 0.4 | 0.4 | 0.4 | 0.4 | 0.4 | 0.4 | 0.4 | 0.4 | 0.4 | 0.4 | 0.4 | 0.4 | 0.4 | 0.4 | 0.4 | 0.4 | 0.4 | 0.4 | 0.4 | 0.4 | 0.4 | 0.4 | 0.2 | 0.2 | **0.0** | 0.2 |
| *Sarcophilus harrisi* | 0.4 | 0.4 | 0.4 | 0.4 | 0.4 | 0.4 | 0.4 | 0.4 | 0.4 | 0.4 | 0.4 | 0.4 | 0.4 | 0.4 | 0.4 | 0.4 | 0.4 | 0.4 | 0.4 | 0.4 | 0.4 | 0.4 | 0.2 | 0.2 | 0.2 | **0.0** |

### 3.3 Deamidation of identified collagen peptides

The deamidation ratios of asparagine (N) and glutamine (Q) residues were calculated for each sample by counting the number of observed deamidated N or Q residues in the peptide data and dividing by the total number of observed N or Q residues and multiplying by 100 to calculate a percentage of deamidated N or Q residues. This ratio was then subtracted from 100 to calculate the remaining non-deamidated N and Q residues across the samples (Figure 6). The deamidation ratios for the AC, AI, and AM datasets of each individual sample were averaged to calculate the standard deviation of the mean. On average, more deamidated N residues were observed than Q residues across samples. The deamidation ratio of Q ranged from ~20% to ~40% across the sample group, while the deamidation ratio of N ranged from ~40% to ~65% across the sample group.

**Figure 6:**
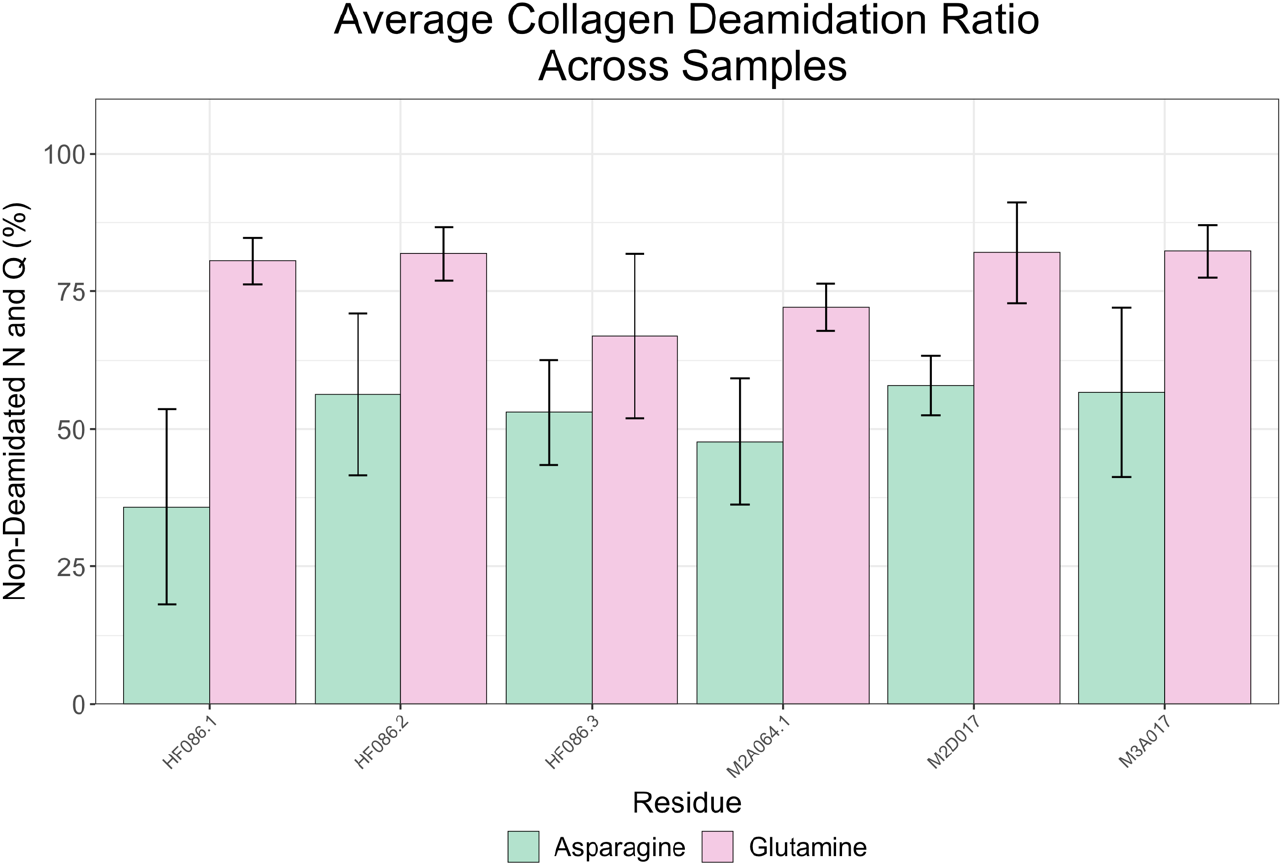
Average asparagine (green) and glutamine (pink) deamidation of collagen peptides identified in the bone artefacts. Error bars denote one standard deviation. Deamidation calculations performed from GPM peptide outputs in R (version 3.6.3) as previously described (Multari, et al., 2022).

## 4. Discussion

The worked bone artefacts analysed in this study were among a few thousand bone fragments excavated from the 63, 65A & 69-71 Harris Street and 14-16 Mount Street, Pyrmont site, and were the only bone pieces to display signs of being worked. The distribution of morphologically identified taxa in the bone assemblage can be seen in Figure 7. The most predominant taxa identified in this assemblage were caprids (Goat, *Capra hircus*; and Sheep, *Ovis aries*), followed by several species of fish, and rodents (Mouse, *Mus musculus*; and Rat, *Rattus norvegicus*). Contextually, the high abundance of caprids and fish in comparison to other taxa in this assemblage is consistent with known records of sheep and goat husbandry, and with the substantial fishing industry that was active in the area at the time (Coghlan, 1969). Meat became a staple in the diets of all social strata in Australia during the 19^th^ Century, because of its wide availability and affordability (Davison, 1987). Almost 10% of the identified bones at the 63, 65A & 69-71 Harris Street and 14-16 Mount Street, Pyrmont site displayed signs of butchery and gnawing, particularly those belonging to caprids, cows, pigs, rabbits, and poultry.

**Figure 7:**
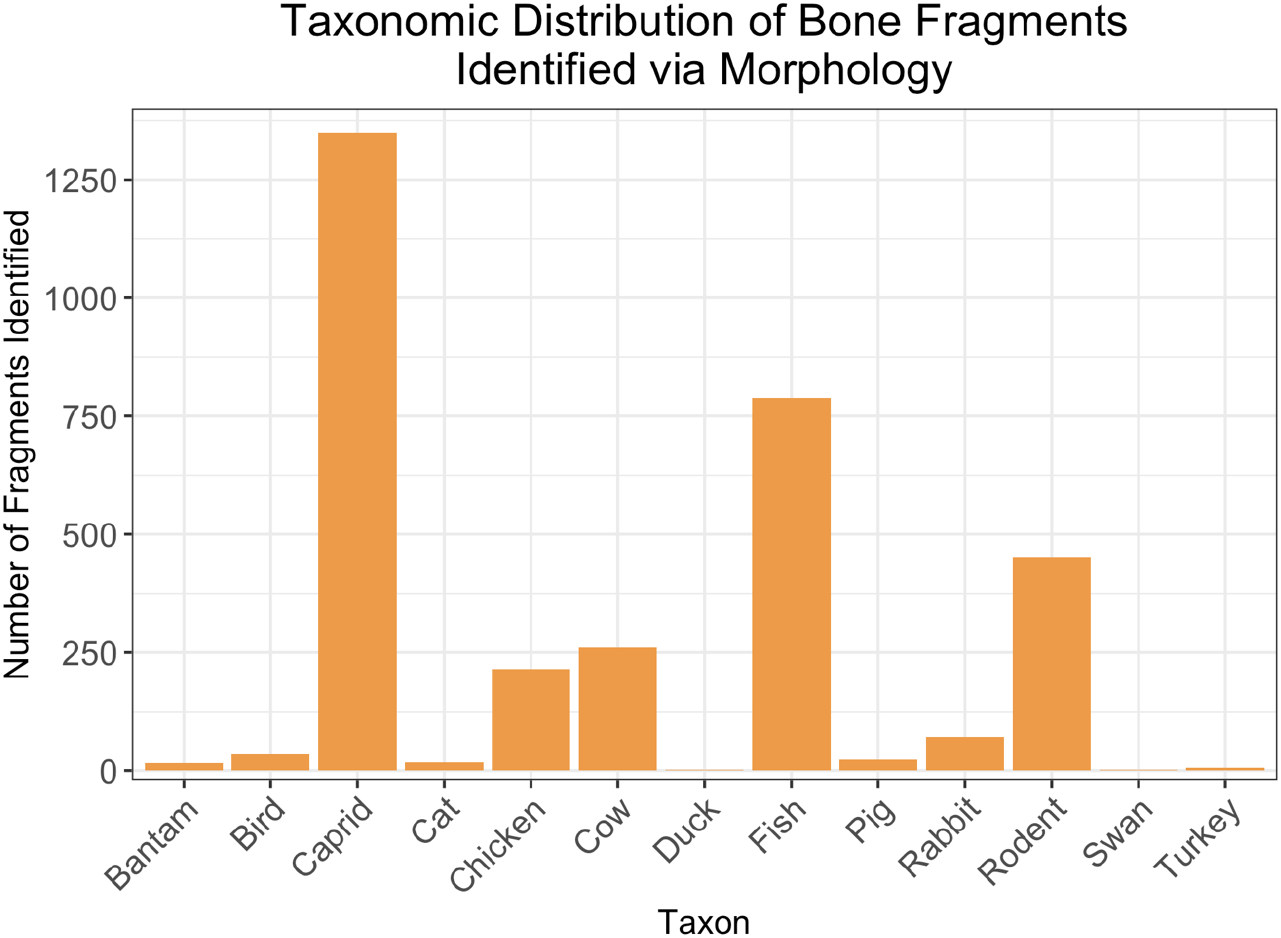
Taxonomic Distribution of bone fragments within the 63, 65A & 69-71 Harris Street and 14-16 Mount Street, Pyrmont assemblage. Count based on number of bone fragments positively identifiable using morphological analyses (Performed by MH in 2017 as part of initial post-excavation analysis). Taxa which are difficult to morphologically discern were grouped accordingly: Caprid includes Goat (*Capra hircus*) and Sheep (*Ovis aries*), and Rodent includes Mouse (*Mus musculus*) and Rat (*Rattus norvegicus*). Figure was produced using R (version 3.6.3) in RStudio (version 1.3.1073) (R Core Team, 2020).

All six of the worked bone artefacts analysed in this study returned highly confident positive identifications for *B. taurus* COL1A1 and COL1A2 peptide markers. Due to the existence of more diagnostic peptide markers, COL1A2 is seen as a more reliable target for the species identification of bone, as seven of the nine standardised peptide regions originate from the COL1A2 protein sequence (Brown, et al., 2021). Table 2 highlights that the collective list of peptide marker sequences identified in each bone handle correlate directly to the COL1A2 sequence for *Bos taurus* and show no direct matches to other species. The identification of these artefacts as cow bones is interesting as cows were not highly represented in the bone assemblage excavated from this site (Figure 7), and the cattle industry was not extensively developed in this region during the early to mid-19^th^ Century (Coghlan, 1969, Blake, 2010). Cattle rearing in 19^th^ Century Australia was considered costly as cattle farmers had to compete with the successful sheep farming industry. Cattle by-products, such as hide, tallow, and dairy, were considered secondary industries and were not nearly as profitable as wool, which meant the price of beef was highly dependent on the success of the wool and mutton industries (Coghlan, 1969, Blake, 2010). As a result, it is likely that abattoirs had to utilise every component of the cow, including the bones, to make a profit. The size, shape, and strength of cattle long bones makes them ideal candidates for knife handle manufacturing. Another plausible explanation for the use of cattle bones to make these artefacts stems from the socio-economic and industrial status of Pyrmont residents and the industrial presence in the area during this period. Pyrmont in the 19^th^ Century was a working-class district occupied by residents of lower socio-economic status who were often too poor to regularly afford expensive meats such as beef over more readily accessible mutton or goat (Blake, 2010). Additionally, such poverty resulted in a necessary reliance in resourcefulness and recycling of materials as the luxury of buying new when something broke was simply not feasible (Ricardi, 2019). The region was home to industries such as abattoirs, butcheries, and fisheries, which could suggest scrap bone was readily available in this area for re-use in repairing small household objects such as cutlery (Coghlan, 1969, Ricardi, 2019). While these bone artefacts were excavated in Australia, it is possible that they may have been manufactured overseas and imported at any point in the colonial history of the region before their final deposition in the mid-19^th^ Century. Stable isotope analysis of ^87^Sr/^86^Sr would potentially provide an answer to this question (Giovas, et al., 2016). However, it would still not be possible to determine whether the bones were imported into Australia in a live animal or as worked artefacts.

Interestingly, the α2 757 marker peptide was not observed in any sample across all three collagen extraction methods. In the *B. taurus* COL1A2 sequence, this peptide marker has a m/z value of either 3017 (with 4 observed hydroxyprolines) or 3033 (with 5 hydroxyprolines) (Brown, et al., 2021). It is likely this peptide was not identified due to incomplete trypsin digestion resulting from the complex helical nature of collagen proteins, as peptides of larger m/z values and sequence length were identified in the datasets. Figure 5 illustrates that no one collagen extraction method was consistently better at recovering peptide marker regions, although the minimally-destructive AM protocol was able to recover at least 1 peptide for each marker region across all six samples. Based on our data, we suggest, wherever feasible, that all three approaches be utilised to maximise peptide recovery and increase confidence in subsequent species identification.

Deamidation has been reported to show promise in being a molecular marker of age in ancient protein studies and is frequently used to separate contaminant proteins from endogenous data (Ramsøe, et al., 2020, Chowdhury and Buckley, 2021). As proteins degrade, the amide bonds in N and Q amino acids undergo a hydrolysis reaction wherein a mass shift of 0.984 Da is observed per deamidation site (where the -NH2 group on the amino acid sidechain is replaced with an -OH) (Jin, et al., 2022). Analysis of the degree of deamidation identified in the collagen peptides showed that the peptides were not heavily deamidated with N residues averaging approximately 40 – 60% non-deamidated and Q residues averaging approximately 60 – 80% non-deamidated across the bone handles (Figure 6). These values are not demonstrably different to modern (<50 years old) reference samples previously analysed in our lab (data not shown). This is likely due to the handles being relatively modern (~170 years old) when compared to more ancient samples such as bone collagen from the Pleistocene which have been used in previous studies to demonstrate significant differences in deamidation status between modern and ancient bones (Ramsøe, et al., 2020). Furthermore, the reliability of deamidation as an index of age versus sample preservation is an active area of debate in the field of paleoproteomics (Schroeter and Cleland, 2016, Chowdhury and Buckley, 2021). More research is required in this field to fully understand the various factors (such as protein structure, sample type, predepositional influences, and the postdepositional environmental conditions) that may be involved in the rate at which deamidation occurs under different conditions (Chowdhury and Buckley, 2021, Hendy, 2021).

## 5. Conclusion

We were able to positively identify as cow bones the material used to create a series of six early colonial Australian worked bone artefacts, using an LC-MS/MS-based ZooMS workflow. All three collagen extraction methods were able to generate peptides compatible with species identification, and it was found that a combinatorial approach is best, wherever feasible. The successful identification of these artefacts as being made from bovine bones has numerous implications on our knowledge of life in the early European settlement of Australia regarding resource availability, animal husbandry and use, and potential trade.

## Abbreviations

ACN: Acetonitrile
NH_4_HCO_3_; AmBiC: Ammonium Bicarbonate
FA: Formic acid
MALDI-ToF/ToF: Matrix-assisted Laser Desorption/Ionisation – Tandem Time of Flight mass spectrometry
MS: Mass Spectrometry
NanoLC-MS/MS: Nanoflow high-pressure liquid chromatography – tandem mass spectrometry
TFA: Trifluoroacetic acid
ZooMS: Zooarchaeology by Mass Spectrometry

## Acknowledgements

DHM acknowledges scholarship support from Macquarie University. Some of the research described herein was facilitated by access to the Australian Proteome Analysis Facility (APAF) funded under the Australian Government’s National Collaborative Research Infrastructure Strategy (NCRIS)/Education Investment Fund. We would like to thank Dr. Belinda Schiller, Dr. Luke Carroll, Dr. Gene Hart-Smith, and Dr. Ardeshir Amirkhani from APAF for their assistance. Special thanks to Patrick da Roza for assistance with the PRIDE repository upload. We thank Michael Rampe for photographing the bone artefacts for this paper. PAH thanks Michael Shaw for continued support and encouragement. The authors gratefully acknowledge Pamela Kottaras from EMM Consulting for providing access to excavation reports, materials, and figures. The excavation from which the bone artefacts were acquired was completed by EMM Consulting in 2017 in compliance with City of Sydney approval (D/2016/916 and D/2017/464) and approval under section 140 of the *Heritage Act 1977* 2017/s140/05.

